# Intracranial hypertension drives astrocyte-mediated neuroinflammation through Piezo1-dependent EGFR activation

**DOI:** 10.1101/2025.05.02.651431

**Authors:** Zongren Zhao, Andrea Hoffmann, Fan Sun, Tamara Merz, Florian Olde Heuvel, Burak Özkan, Franziska Münz, Enrico Calzia, Michael Gröger, Sandra Kress, Peter Radermacher, Francesco Roselli, Thomas Kapapa, Marica Pagliarini

## Abstract

Intracranial hypertension is a major driver of secondary injury after acute subdural hematoma (ASDH), yet how mechanical stress is translated into neuroinflammatory signaling remains poorly understood. Here, we identify a mechanosensitive astrocyte signaling pathway that links elevated intracranial pressure (ICP) to inflammatory amplification in the injured brain.

Using a clinically relevant porcine ASDH model combined with mechanistic studies in human iPSC-derived astrocytes, we demonstrate that sustained ICP elevation induces bilateral neuroinflammation together with coordinated upregulation of mechanosensitive ion channels and receptor tyrosine kinase (RTK) pathways. Integrative analysis of molecular and physiological datasets identified astrocytes as the principal cellular responders to ICP and revealed epidermal growth factor receptor (EGFR) as the astrocyte-associated RTK most strongly correlated with ICP dynamics, inflammatory chemokine expression, and survival. Pharmacological activation of the mechanosensitive channel Piezo1 in human astrocytes was sufficient to trigger EGFR internalization, site-specific phosphorylation, and ERK signaling, promoting structural remodeling and robust induction of pro-inflammatory mediators including CCL2, IL-6, and IL-8. Conversely, EGFR inhibition attenuated inflammatory signaling while enhancing astrocytic programs associated with water handling and edema containment. In vivo, increased expression of EGFR ligands together with elevated EGFR phosphorylation supported sustained pathway engagement following ASDH, and correlation analyses linked Piezo1 expression and EGFR activation with ICP severity and adverse outcome.

Together, these findings define a mechanotransduction axis in which astrocytic Piezo1 signaling integrates mechanical stress with EGFR-dependent neuroimmune responses, positioning EGFR as a translationally accessible target to modulate inflammation-driven secondary brain injury.

## Introduction

Intracranial hypertension (ICH), defined as the increase of IntraCranial Pressure (ICP) above 15-20mmHg, is a life-threatening phenomenon associated with traumatic and cerebrovascular brain lesion [1]., ICH is associated not only with intracranial volume-occupying lesions such as acute subdural haematoma (ASDH) or post-traumatic parenchymal necrotic/haemorrhagic lesions [2], but also with the brain edema that accompanies these conditions [3]. Unchecked increase in ICP paves the way to brain herniation and to diffuse cerebral hypoxia-hischemia, with substantial acute mortality and unfavourable long-term neurological prognosis [4, 5].

Despite its relevance, treatments addressing ICP are currently limited to hyperosmolar therapy, hyperventilation and eventually surgical decompression [6]. Of note, in ASDH increased ICP may persist even after the original haematoma has been drained [7], indicating an ongoing pathogenic process driving ICH. None of the currently available options addresses the pathophysiology underlying the increased ICP.

Although it is conceivable that the initial response to intracranial volume-occupying space such as the haematoma in ASDH, and ICH may be triggered by the mechanical stress imposed on brain parenchyma, meninges and vascular structures [8, 9], the number of pathways involved in secondary ICH, i.e. not related to the initial mechanical lesion, is very large and include hypoxia/ischemia [7, 10], iron toxicity [11], vascular and cytotoxic edema [12] and neuroinflammation.

Ideally, identifying the upstream drivers of the response to ICP raise that make ICH a persistent phenomenon in conditions such as ASDH, before the extensive diversification and divergence of the pathophysiological cascades, would allow an early and effective treatment of the ICH. Receptor Tyrosine Kinases (RTK) constitute ideal candidates for this role, since RTK are highly expressed in multiple structures and cellular subpopulations of the cerebral parenchyma [13, 14], provide entry points to multiple core cellular functions (such as proliferation, inflammation and apoptosis;[15, 16] and are extensively activated upon injury [17]. Furthermore, RTKs are targeted by a growing number of inhibitors developed for cancer treatment and already approved for human use [18], which lend themselves to drug repurposing.

We elected to investigate the dysregulation of RTK and their mechanistic involvement in the response to raised ICP occurring upon ASDH, exploiting a porcine model, which offers not only the advantage to be established in a gyrencephalic animal (in contrast to more diffuse murine models;[19]) but also to provide a clinically realistic physiological monitoring and a human-rated neurointensive care condition [20] as well as a characterized pathology [20-22]. We have paired the in-vivo exploration with in vitro human IPSC-derived astrocytes to address mechanisms at molecular level and highlight the relevance of the findings for human patients.

## Material and methods

### Animal model and surgery

All experiments were conducted in accordance with the National Institutes of Health guidelines and the European Directive 2010/63/EU and were approved by the local Animal Care Committee and the Federal Authorities for Animal Research (Regierungspräsidium Tübingen; Reg.-Nr. 1559).

The porcine model of ASDH has been previously reported and characterized and the detailed information are reported in the Supplementary Methods [20, 23]. Briefly, for the instrumentation and the generation of the experimental ASDH, animals were premedicated with intramuscular azaperone (5 mg/kg) and midazolam (1–2 mg/kg). Anesthesia was induced with intravenous propofol (1–2 mg/kg) and ketamine (1 mg/kg), followed by endotracheal intubation and controlled mechanical ventilation. Anesthesia was maintained with continuous propofol and remifentanil infusions. Ventilation was adjusted to maintain normocapnia (PaCO₂ 35–40 mmHg). Maintenance fluids consisted of balanced electrolyte solution (10 mL/kg/h). Femoral arterial and venous access was established for continuous hemodynamic monitoring and blood sampling. A PiCCO catheter was used for advanced cardiovascular monitoring. Burr holes were drilled bilaterally, and subdural drains were placed. A multimodal intraparenchymal probe was inserted subcortically for continuous monitoring of intracranial pressure (ICP), brain temperature, and brain tissue oxygenation. Hemorrhagic shock was induced by controlled passive blood withdrawal (≈30% blood volume), while maintaining cerebral perfusion pressure (CPP) ≥50 mmHg. After a 60-minute stabilization period, ASDH was induced by subdural injection of autologous blood (0.1 mL/kg) over 15 minutes. This subdural blood volume was injected because 10 % of the intracranial volume represents the threshold for supra-tentorial volume tolerance [24]. Physiological parameters, including ICP, CPP, brain oxygenation, arterial pressure, and oxygen saturation, were continuously monitored. Two hours post-injury, targeted resuscitation was initiated, including retransfusion of shed blood, fluid administration, and norepinephrine titrated to restore mean arterial pressure and CPP to baseline values (≥60 mmHg). Neurological status was assessed daily using a modified Glasgow Coma Scale for pigs. Animals were euthanized upon predefined humane endpoints (CPP below 60 mmHg despite receiving the maximum dose of vasopressors, with a heart rate limit of 160 beats per minute to prevent myocardial injury from tachycardia and/or if they developed acute anuric kidney failure with hyperkalemia (blood potassium > 6 mmol/l) and cardiac arrhythmias). Naïve control animals (n = 6) underwent anesthesia alone without any neurosurgical instrumentation and induction of ASDH and hemorrhage. following the completion of the surgical catheter placements, adjustments were made to the ventilator settings to achieve an inspiratory/expiratory (I/E) ratio of 1:2, F_i_O_2_ set at 0.21, and zero end-expiratory pressure (0 cmH2O) in order to closely replicate physiological conditions. These control animals were euthanized after a total experiment duration of approximately 2 hours and 15 minutes [25].

### Tissue sampling, RNA extraction and qPCR

After the extraction of the brain and the gross sectioning (approx 1cm) along the rostro-caudal axis, cerebral cortex samples were collected in correspondence of the parietal lobe and from an area adjacent to, but macroscopically free of, the hematoma, and from matching controlateral region. Tissue samples were snap-frozen on dry-ice and stored at -80 until processed. The samples were homogenized in QIAzol reagent (Qiagen Cat nr79306), and total RNA was extracted using a column-based purification method with on-column DNase digestion according to the manufacturer’s instructions. Purified RNA was eluted in RNase-free water, quantified using Nanodrop 1000 and stored at −80 °C until analysis. Species-specific primers for porcine genes were designed using NCBI Primer-BLAST, validated for specificity and efficiency, and are listed in Supplementary Table 1. Quantitative PCR was performed using SYBR Green chemistry on a QuantStudio 1 system. Reactions were run in duplicate, GAPDH was used as the housekeeping gene, and relative mRNA expression levels were calculated using the 2^−ΔCt method as previously reported [26].

### Protein extraction and Western Blot

Protein concentrations were determined using the BCA Protein Assay Kit (Thermo Fisher Scientific Cat. Nr. 232225) according to the manufacturer’s instructions. A total of 75 µg of protein was mixed with 1× loading buffer, 5% 2-mercaptoethanol, and RIPA buffer, then heated at 95°C for 5 minutes.

Protein samples were resolved on an 8 or 12% SDS-PAGE gel and subsequently transferred to a polyvinylidene difluoride (PVDF) membrane for 2h at room temperature. The membranes were blocked with 5% bovine serum albumin in 1× Tris-buffered saline with Tween-20 (TBST) for 1 hour at room temperature.

Following blocking, the membranes were incubated with primary antibodies on a shaker at 4°C overnight. After three washes with 1× TBST for 10 minutes each at room temperature, membranes were incubated with secondary antibodies diluted in 5% BSA in 1× TBST for 2 hours at room temperature on a shaker. The membranes were then washed three times for 10 minutes each with 1× TBST at room temperature. The specifications of the antibodies used are reported in Supplementary Table 2.

Protein detection was performed using an enhanced chemiluminescence detection system (Biorad, Cat nr.1705061). Band intensity was quantified using ImageJ software, with normalization to the β-actin loading control.

### Differentiation of human iPSC-derived NPCs into astrocytes

Human iPSC-derived neural progenitor cells (STEMCELL Technologies, SCTi003-A) were expanded according to the manufacturer’s protocol and differentiated into astrocytes between passages P3–P5 following progenitor expansion and splitting. Cells were plated on Matrigel-coated cultureware (Corning, Cat. No. 354277) at a seeding density of approximately 1–1.5 × 10⁵ cells/cm². Astrocyte differentiation was performed using DMEM (Thermo Fisher Scientific) supplemented with N-2 Supplement (1%; Thermo Fisher Scientific), GlutaMAX Supplement (2 mM; Thermo Fisher Scientific), and fetal bovine serum (1%; Thermo Fisher Scientific), Penicillin/Streptomycin (1%, Thermo Fisher Scientific). Astrocyte differentiation medium was stored protected from light at 2–8 °C and used within 4 weeks, with regular medium changes performed according to the manufacturer’s instructions. Astrocytic maturation was confirmed by quantitative PCR analysis of S100β expression.

### Astrocytes treatment

Astrocytes were seeded and let grow until reaching 70-80% confluency. One day prior to stimulation, the culture medium was replaced with serum-free medium and cells were incubated overnight. On the following day, cells were divided into the following groups: Control; EGF (10 ng/mL, 2 hours, 48h); Yoda (2 μM, 2 hours); and Gefitinib (100 nM, 30 minutes) followed by Yoda (2 μM, 2 hours), ERK inhibitor PD98059 (20 µM 30 minutes) followed by either EGF (10 ng/ml 2h) or Yoda (2 µM 2 hours). After treatment, the culture medium was removed, and cells were washed with PBS and collected for subsequent RNA and protein extraction.

### RNA extraction from human astrocytes and qPCR

Human iPSC-derived astrocytes cultured in vitro were washed with cold PBS and lysed directly in the culture wells using QIAzol reagent according to the manufacturer’s instructions. Total RNA was extracted using the same column-based purification protocol described for porcine cortical tissue. Purified RNA was eluted in RNase-free water, quantified using a Nanodrop spectrophotometer, and stored at −80 °C until further analysis. cDNA synthesis and quantitative PCR were performed using SYBR Green chemistry on a QuantStudio 1 system following the same amplification conditions described above. Species-specific primers targeting human transcripts were designed using NCBI Primer-BLAST. GAPDH was used as the housekeeping gene, and relative mRNA expression levels were calculated using the 2^−ΔCt method.

### Astrocyte proteins extraction and EGFR phosphorylation array

Astrocytes cultured in 6-well plates were washed with cold PBS and lysed directly in the well using 500 µL of RIPA buffer supplemented with protease and phosphatase inhibitors (Roche). Lysates were incubated on ice for 20–30 min, sonicated three times for 3 s each, and clarified by centrifugation. Protein concentration was determined, and 250 µg of total protein per sample was used for analysis. EGFR phosphorylation was assessed using a human phospho-EGFR antibody array performed according to the manufacturer’s instructions. Briefly, array membranes were blocked for 1 h, incubated with diluted cell lysates overnight at 4°C, washed, and incubated with a biotin-conjugated anti-EGFR antibody cocktail followed by HRP-conjugated streptavidin. Signals were detected by chemiluminescence and imaged with a 16 bit resolution using a digital imaging system (LAS 4000).

### Astrocytes live-cell holotomographic imaging

Holotomographic live-cell imaging was performed using a holotomographic microscope (Nanolive) to obtain high-contrast, label-free imaging of human iPSC-derived astrocytes. Cells were imaged under physiological conditions (37 °C, 95/5% O₂/CO₂) using a 60× air objective (NA 0.8). Three-dimensional holotomographic stacks were acquired at defined intervals for a total duration of 2 h following treatment. Astrocytes were exposed to vehicle, EGF, Yoda-1, or Yoda-1 in combination with the EGFR inhibitor gefitinib, added immediately prior to imaging. Cells were cultured on optical-grade 35 mm dishes, and image acquisition was performed without fluorescent labeling. Image datasets were exported using the STEVE software suite (Nanolive). Quantitative analysis of time-lapse holotomographic image sequences was performed using ImageJ. Three-dimensional tomographic datasets were reconstructed and exported as time-series images for analysis. Cell boundaries were segmented based on refractive index contrast to generate binary masks defining individual astrocyte areas. Segmentation parameters were kept constant across all experimental conditions, and masks were visually inspected to ensure accurate identification of cell borders. Cell area was calculated from the segmented regions using standard ImageJ measurement tools. Endoplasmic reticulum structures were identified and segmented based on their characteristic refractive index distribution within the holotomographic images, allowing label-free discrimination from surrounding cytoplasmic regions. Cell motility was quantified using the TrackMate plugin in ImageJ by tracking the centroid position of individual cells across sequential time frames. Net displacement and total displacement were calculated from centroid trajectories as measures of cell movement, using identical tracking parameters for all experimental groups.

### Immunocytochemistry of hIPSC derived astrocytes

After treatment, the coverslips were washed two times with ice cold DPBS-/- and fixed for 15 min with ice cold 4% PFA added with 0.1 M sucrose. The cells were washed 2 times with ice cold DPBS-/- and blocked in PBS-/- with 3% BSA and 0.3% Triton-X 100 at RT for 2 h. Subsequently, cells were incubated with primary antibodies (S100b 1:300 Sysy Cat nr 287 004, EGFR 1:200 CST Cat nr 4267S) diluted in a blocking buffer at 4°C for 48 h. After washing with PBS-/- three times for 30 min, cells were incubated with secondary antibodies at RT for 2 h and washed three times for 30 min with PBS-/- at RT. Finally, the coverslips were mounted on microscope glass slides with Fluorogold prolong antifade mounting medium (Invitrogen).

### Bioinformatic analysis

Cluster analysis was conducted using the K-means algorithm to identify patterns of correlation among molecular markers in scikit-learn module of Python 3.11.6. To visualize the results was used matplotlib [27]. The number of cluster was determined by a distance threshold: threshold ≈ 0.7 × max linkage height. To estimate the relative contributions of individual cell types to the gene expression profile of the most prominent cluster, we generated a cell type–specific gene expression signature matrix using single-cell RNA sequencing data annotated with cell types from the Brain RNA-seq database [28]. For each annotated cell type, we computed the average gene expression across all cells, resulting in a signature matrix with genes as rows and cell types as columns.

### Statistical analysis

Normality was assessed for all groups using the Shapiro–Wilk test. Correlations were evaluated using Pearson’s correlation coefficient (r) with corresponding p-values. Quantitative data are presented as mean ± standard deviation (SD). For comparisons between two groups, an unpaired t-test was used for normally distributed data, and the Mann–Whitney U test was applied when normality was not met. For comparisons involving three or more groups, one-way ANOVA with Bonferroni post-hoc was performed. A p-value < 0.05 was considered statistically significant. All analyses were conducted using GraphPad Prism version 9.0 (GraphPad Software).

## Results

### Intracranial hypertension and bilateral cerebral cortex involvement in the porcine Acute Subdural Hematoma (ASDH) model

The subdural injection of 0.1ml/kg of autologous blood in correspondence of the parietal lobe (Fig. 1A) resulted in the formation of a subdural haematoma, whose location and extension were verified by necropsy in all cases; some blood percolation or diffusion to subarachnoid spaces was sporadically observed (1/10; Fig. 1B). Coherent with the appearance of the acute subdural haematoma, increased in ICP measured in both hemispheres, and decreased CPP were detected (with reference to previously reported physiological ranges: ICP 5-10 mmHg, CPP 110-125 mmHg [20]) soon after the intracranial injection (Fig. 1C). The animals (n=10) were sacrificed between 36 and 48h after the intracranial blood injection, according to pre-specified termination criteria (see methods).

**Figure 1.**
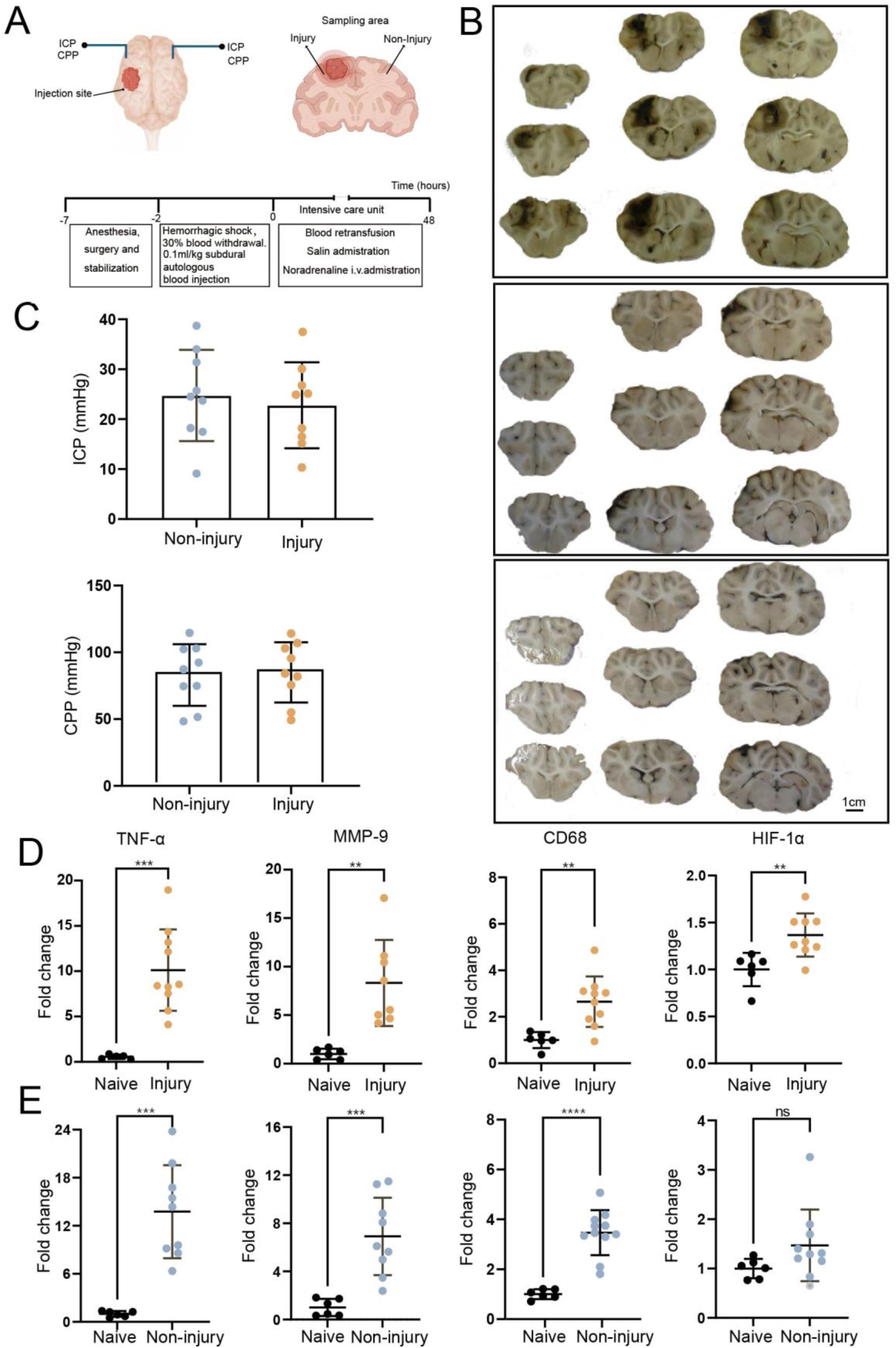
Acute subdural hematoma induces intracranial hypertension and bilateral cerebral tissue involvement in a porcine model. (A) Experimental design of the acute subdural hematoma (ASDH) model, including subdural injection of autologous blood and physiological monitoring (ICP,CPP). (B) Representative macroscopic brain sections obtained at necropsy showing subdural hematoma predominantly in the ipsilateral hemisphere, with variable extension into the subarachnoid space and brain parenchyma. Occasional bilateral involvement was observed. Scale bar: 1 cm. (C) Intracranial pressure (ICP) and cerebral perfusion pressure (CPP) following ASDH induction, demonstrating elevated ICP and reduced CPP in injured animals in ipsilateral and contralateral side compared to reference range. (D–E) Relative mRNA expression levels of TNF-α, MMP-9, CD68, and HIF-1α in cortical tissue collected at the end of the observation period (36–48 h). TNF-α, MMP-9, and CD68 were increased bilaterally, whereas HIF-1α was significantly upregulated in the ipsilateral hemisphere but not in the contralateral hemisphere. Data are presented as fold change to naïve controls. Unpaired t-test was performed. Naïve animals n = 6; ASDH animals n = 10. *p < 0.05, **p < 0.01, ***p < 0.001.

For validation of the samples, we monitored the expression of a set of genes marker for hypoxia, neuroinflammation and tissue damage, contrasting cortical samples from injury-naive animals (subject to anesthesia but not to instrumentation or surgery; n=6) to ASDH animals (cortical samples collected either ipsilateral either contralateral to the site of blood injection). Expression of TNF-α and of the phagocytes markers CD68 and MMP-9 was upregulated both in ipsi- and controlateral samples (Fig. 1D-E). On the other hand, expression of the hypoxia-inducible factor 1-alpha (HIF1-α) was upregulated only in the samples ipsilateral to the subdural haematoma, indicating regional hypoperfusion/hypoxia (Fig. 1D-E). Thus, the ASDH results in ICP increase, ipsilateral hypoxia and bilateral neuroinflammation.

### ASDH drives the upregulation of multiple mechanosensitive channels in the porcine cortex

ASDH generates a pronounced mass effect that deforms and compresses the underlying (and, due to the relative flexibility of the falx, also the contralateral) cerebral tissue. We hypothesized that the mass-effect and the diffuse mechanical strain (alone or together with the local hypoxia/hypoperfusion) may modify the expression of mechanosensitive ion channels.

We prioritized a list of mechanosensitive targets from the Piezo (Piezo1, Piezo2) and from the TRCP (TRCP1-7) families with known expression in the brain and we explored their mRNA levels in cortical samples from ipsilateral and contralateral hemispheres as well as in cortical samples from naive animals. We observed a robust upregulation of both Piezo1 and Piezo2 (Fig. 2A-B), in the ipsi- and in the contralateral samples. Likewise, TRPC1, TRPC3, and TRPC4 were upregulated bilaterally (Fig. 2C-E), whereas TRPC6 and TRPC7 were selectively upregulated only in the ipsilateral hemisphere (Fig. 2G, H). TRPC5 expression remained unchanged (Fig. 2F).

**Figure 2.**
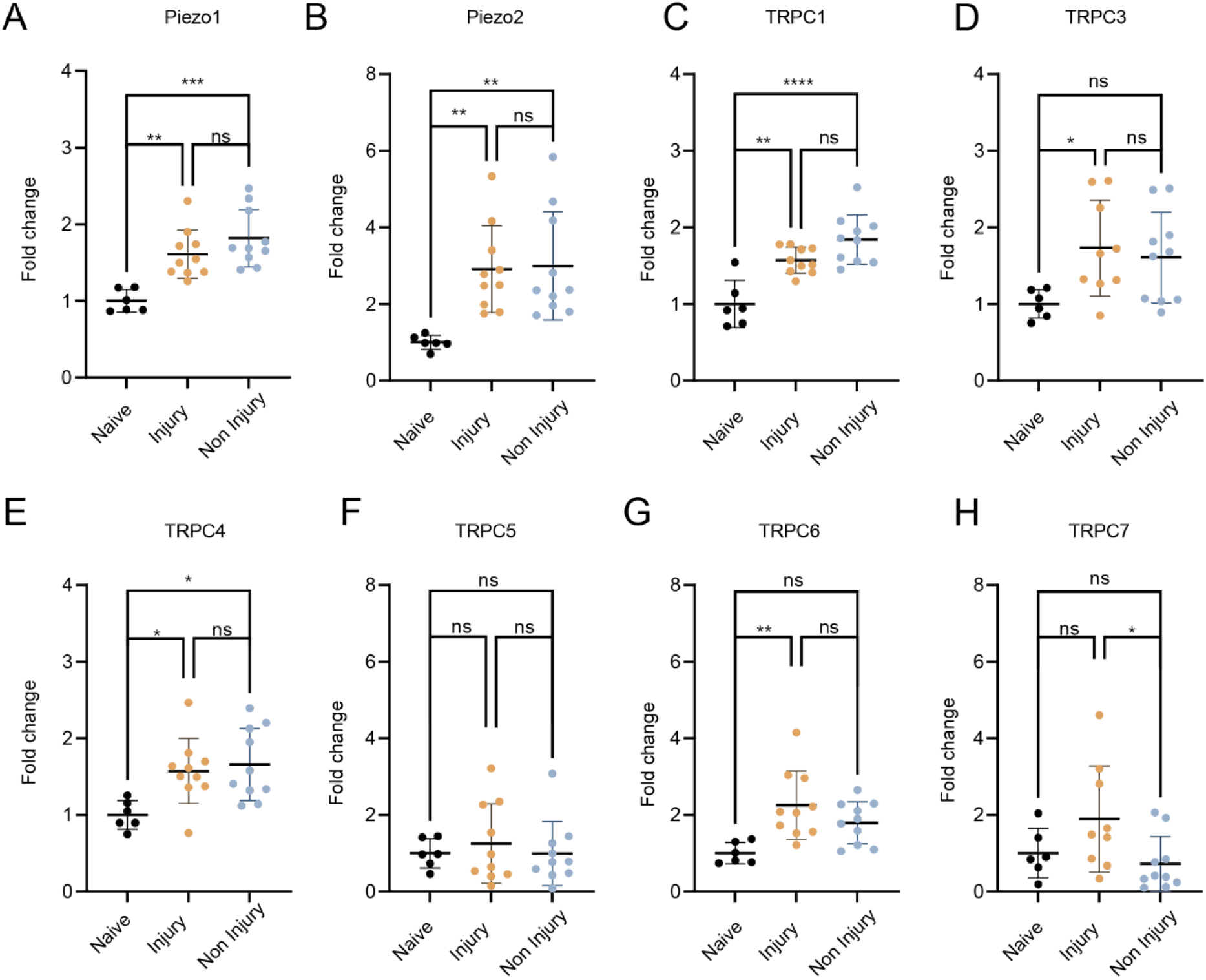
Acute subdural hematoma induces bilateral upregulation of mechanosensitive ion channels in the porcine brain. (A–H) Relative mRNA expression levels of Piezo1 (A), Piezo2 (B), TRPC1 (C), TRPC3 (D), TRPC4 (E), TRPC5 (F), TRPC6 (G), and TRPC7 (H) in naïve control, ipsilateral injury, and contralateral non-injury hemispheres following ASDH. The data are presented as fold change relative to naïve controls. Each dot represents an individual animal; bars indicate mean ± SD. Statistical analysis: One way ANOVA with Tuckey post-hoc (ns, not significant; *p < 0.05; **p < 0.01; ***p < 0.001; ****p < 0.0001). Naive animals n=6, ASDH (injury side and non-injury side) animals n=10.

Together, these findings demonstrate that ASDH induces an across-the-board upregulation of mechanosensitive ion channels taking place in both hemispheres.

### Intracranial hypertension drives an astrocyte-specific transcriptional response upon ASDH

Having established the transcriptional footprint of ASDH on mechanosensors, we set out to determine the identity of the cellular subpopulations responding to ICP with such upregulation profile, and the cell-specific pathways set in motion by increase in ICP that follows ASDH, focusing on the ipsilateral cortical samples.

We first quantified the expression of a panel of genes enriched in glia, endothelial, or immune cells (prioritized as the cell types with the highest potential to respond to ICP in ASDH). Astrocyte-associated genes GFAP, AQP4, and S100B were upregulated in the ipsilateral cortex (Fig. 3A). Endothelial and blood–brain barrier–related genes Claudin-5 and VCAM-1 were also significantly increased, whereas ZO-1 expression remained unaffected (Fig. 3B). Likewise, both pro-(IL-1β, CCL2, IFN-γ) and anti-inflammatory (IL-4, IL-10, IL-13) mediators were elevated, consistent with an active neuroimmune response (Fig. 3C).

**Figure 3.**
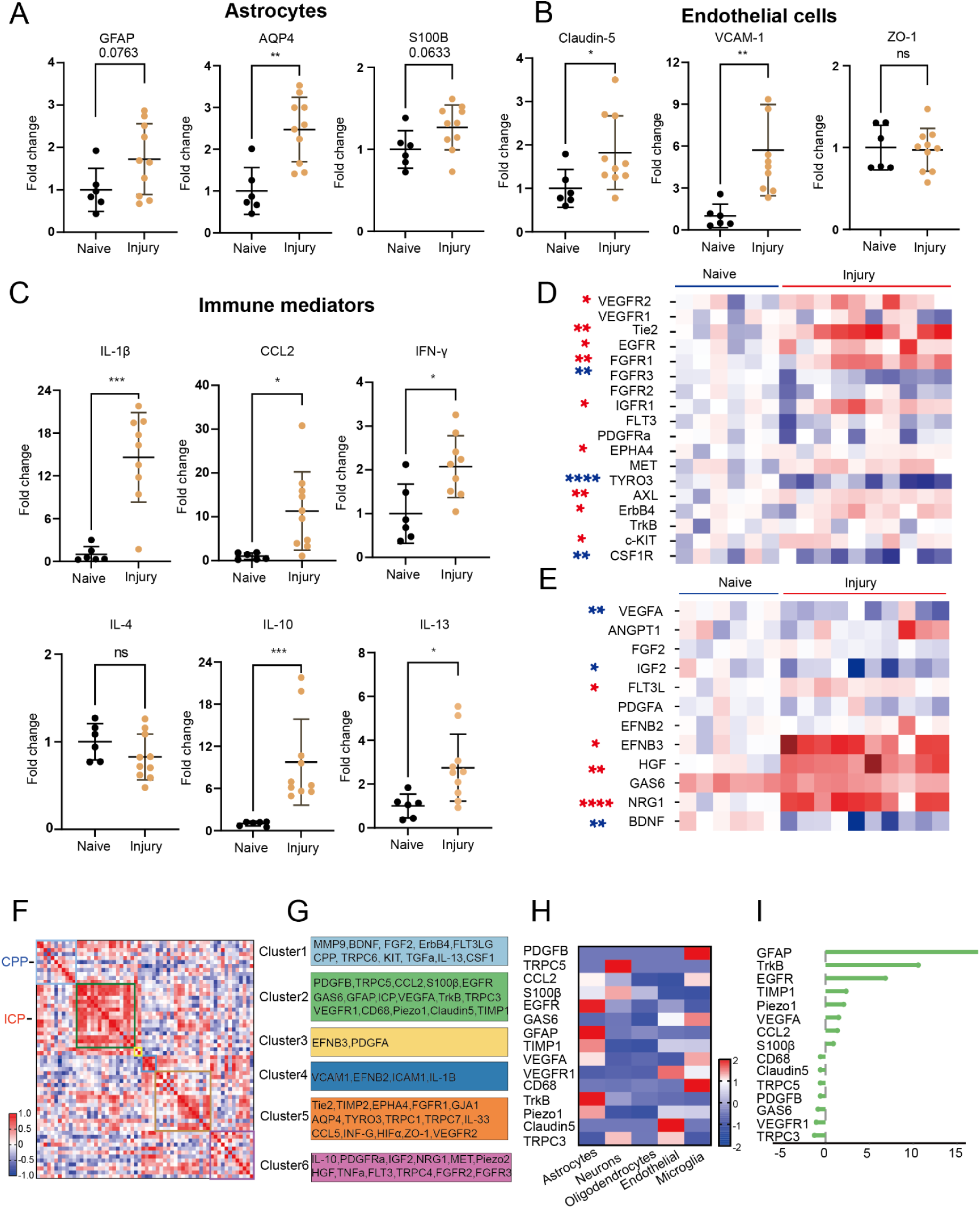
Astrocyte-associated RTK signaling defines an intracranial pressure–linked transcriptional program after ASDH. (A–C) Quantitative PCR analysis of cortical tissue collected 36–48 h after ASDH showing altered expression of astrocytic markers (GFAP, AQP4, S100B), endothelial markers (Claudin-5, VCAM-1, ZO-1), and inflammatory mediators (IL-1β, CCL2, IFN-γ, IL-4, IL-10, IL-13). Data are presented as fold change relative to naïve controls. Naïve animals n = 6; ASDH animals n = 10. * p < 0.05, ** p < 0.01, ***p < 0.001, ****p < 0.0001. (D) Heatmap showing relative expression of selected receptor tyrosine kinases in injured versus naïve cortex. The data are expressed as z score. (E) Heatmap showing relative expression of corresponding ligands associated with the analyzed RTKs. The data are expressed as z score. (F) K-means clustering integrating gene expression data with intracranial pressure (ICP) and cerebral perfusion pressure (CPP), identifying six clusters of co-regulated molecular and physiological variables. (G) Cluster annotation highlighting representative genes within each cluster. (H) Cell-type enrichment analysis across major brain cell populations (astrocytes, neurons, oligodendrocytes, endothelial cells, microglia). (I) Enrichment score ranking indicating preferential association of Cluster 2 genes with astrocytic signatures.

We further explored the cellular specificity and the pathways set in motion by the ASDH by taking into consideration a set of Receptor Tyrosine Kinases (RTK) highly expressed in vascular, glial and immune cells and directly regulating core biological phenomena such as vascular permeability, astrocyte proliferation and inflammatory signaling. Among the 18 RTK, we found that cortex from the ASDH site (compared to naive-animals samples) displayed the significant upregulation of VEGFR1, VEGFR2, Tie-2, EGFR, ErbB4, FGFR1, IGF-1R, HGFR, and Axl, whereas CSF1R and TYRO3 were downregulated (Fig. 3D). We integrated the RTK expression study with the quantification of the mRNA of their cognate ligands, revealing the downregulation of VEGF-A, FGF2, and IGF-1 and the substantial upregulation of Neuregulin-1, HGF, and EphB3 were significantly upregulated, supporting functional engagement of EGFR/ErbB family and Met pathways (Fig. 3E).

In order to identify the cell-types and the pathways most strongly correlated with the ICP, we performed a hierarchical cross-correlation clustering on the gene-expression dataset combined with the ICP and cerebral perfusion pressure (CPP) data. K-means statistics supported the identification of six distinct clusters (Fig. S1; Fig. 3F); interestingly, CPP clustered together with several inflammatory mediators in Cluster1, whereas ICP independently clustered in Cluster2 (Fig. 3G). Cluster 1 included CPP as well as growth factors, extracellular matrix remodeling, myeloid-support signals, and TRPC6, consistent with adaptive tissue remodeling linked to mechanical strain and perfusion changes. Cluster 2 involved ICP as well as several astrocytic markers, chemokines, and mechanosensitive channels (Piezo1, TRPCs), defining a pressure-driven astrocyte reactive program. Of note, among the RTKs, only EGFR, VEGFR1 and TrKB clustered with ICP. Clusters 3 to 6 included several gene sets for vascular, immune and inflammatory responses. Thus, the clustering analysis reveals a differential set of correlations for ICP and CPP and identifies a set of genes closely related to ICP levels (Fig S2).

We further refined our interpretation of CPP and Cluster 1 or ICP and Cluster 2 genes by applying a cell-type annotation based on a single-cell transcriptomics reference atlas [29]. In Cluster 1 (Fig S 2G) the cell deconvolution analysis showed a neuronal enrichment. However, we demonstrated a significant enrichment in astrocytic signatures within Cluster 2 (Fig. 3H–I), suggesting that astrocytes constitute the dominant cellular contributors to the transcriptional program most strongly correlated with ICP. Only limited enrichment for other cell types emerged when neurons, endothelial cells or other glial cells were considered (Fig. S3).

Taken together, the expression data, the clustering and the cell-type deconvolution suggest that astrocytes are the cell type most reactive to ICP and that EGFR, VEGFR1 and TrKB are candidate signaling hubs responding to ICP.

### Piezo1 opener Yoda-1 induces EGFR phosphorylation, signaling and internalization in human astrocytes

We prioritized the study of EGFR signaling in this condition because of its prominent role in regulating astrocyte reactivity and proliferation [30] and its substantial translational outlook (EGFR inhibitors are approved for human use in oncology). Since Piezo1 was among the few mechanosensors co-clustering with ICP as well as with EGFR, we explored the possibility of a mechanistic connection between Piezo1 activation and the upregulation of EGFR. We set out to investigate whether Piezo1 corresponded not only to the increase in EGFR mRNA but also to the activation of the EGFR downstream signaling. We opted to employ human iPSC-derived astrocytes to verify that this pathogenic pathway was conserved between pigs and humans. To simulate the ICP-driven Piezo1 activation we used the Piezo1 opener Yoda-1 [31] and we explored if cultured astrocytes exposed to Yoda-1 displayed the biochemical and morphological hallmarks of active EGFR signaling.

First, to verify that Piezo1 activation resulted in EGFR phosphorylation, we treated human astrocytes with Yoda-1, with EGF (as positive control) or with Yoda-1 together with the EGFR inhibitor Gefitinib and in whole-cell protein extracts we probed the extent of phosphorylation of multiple epitopes on the EGFR intracellular domain using a focused phosphoproteomic array (Fig. 4A-B). EGF treatment (2h) resulted in the significant phosphorylation of all tyrosine and serine epitopes under consideration [31]. Interestingly, Piezo1 activation by Yoda resulted in a more restricted pattern, with a significant increase the phosphorylation of Tyr1068 and Tyr922 (associated with downstream Ras-Erk and Ca²⁺-mediated signaling; [32], of Tyr845 (related to Src signaling; [33]) and of Tyr1045 (involved in receptor internalization and ubiquitination; [34]). Notably, concomitant treatment with Yoda+Gefitinib resulted in the almost complete abolishment of EGFR phosphorylation across all epitopes, indicating that Piezo1-mediated EGFR phosphorylation requires the kinase activity of the EGFR itself (Fig. 4B). We further confirmed the activation of EGFR signaling taking advantage of an antibody directed against the extracellular portion of the EGFR with an antibody-feeding experimental design. Upon treatment of astrocytes with EGF or Yoda, a substantial decrease in the immunolabelling of surface EGFR was detected (Fig. 4C-D), with the corresponding increase in the internalized receptor, a pattern consistent with EGFR autophosphorylation and internalization. Interestingly, EGF, but not Yoda, also induced the clustering of EGFR on the cell membrane, further indicating that Yoda and EGF trigger similar, but not completely overlapping, EGFR phosphorylation patterns.

**Figure 4.**
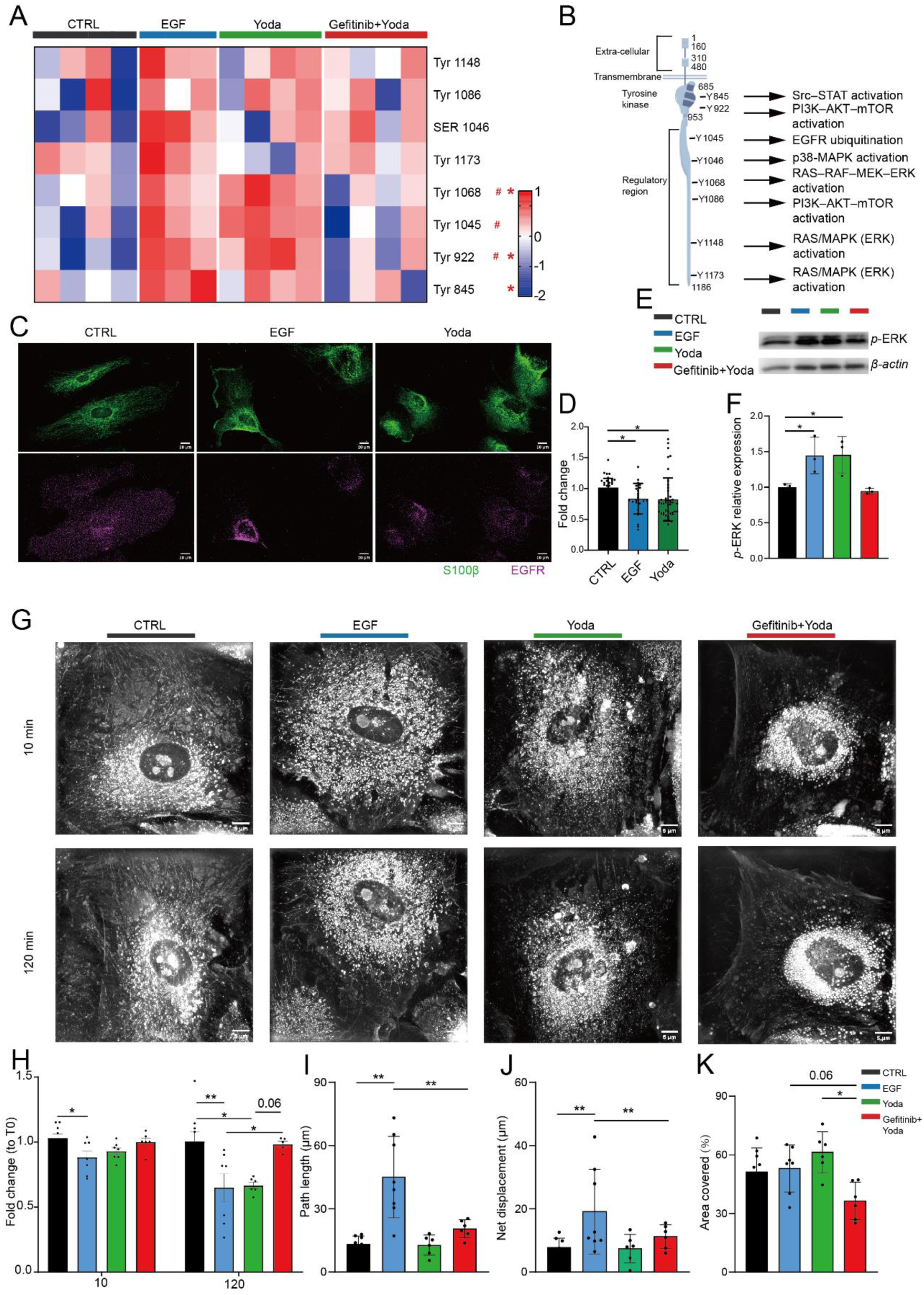
Piezo1 activation induces selective EGFR phosphorylation and structural remodeling in human astrocytes. (A) Phospho-EGFR array heatmap showing site-specific phosphorylation of EGFR residues in CTRL, EGF, Yoda, and Gefitinib+Yoda conditions. (*) indicate significant differences between EGF and CTRL; (#) indicate significant differences between Yoda and CTRL. The data are shown as z-scores (B) Schematic diagram illustrating Yoda-1–induced site-specific EGFR phosphorylation and downstream signaling pathways in human astrocytes. (C) Immunofluorescence images of astrocytes stained for S100β (green) and EGFR (magenta) under CTRL, EGF, and Yoda conditions. Representative images are shown. (D) Quantification of relative EGFR levels across experimental groups. One way ANOVA with Bonferroni post-hoc. * p < 0.05. Three independent experiments were performed. (E-F) Immunoblot analysis and quantification of ERK phosphorylation, demonstrating activation by EGF and Yoda-1 and suppression by Gefitinib. (G) Representative holotomographic images of astrocytes following treatments. H) Cell area measurement showed a reduction overtime. Data are showed as fold change to T0. (I-J) Quantification of astrocyte motility parameters (net and total displacement in µm). (K) Quantification of endoplasmic reticulum size. Data are presented as mean ± SD. One-way Anova with Bonferroni post hoc was used to compare the groups (n=8). Statistical comparisons are shown as follows: * p < 0.05, ** p < 0.01, *** p < 0.001, **** p < 0.0001.

Finally, we assessed if Yoda-1 induced EGFR phosphorylation caused a re-arrangement of cellular architecture (as expected since EGFR signals through rac/cdc42 to the actin cytoskeleton [35]). We used a live-imaging holotomographic system to record changes in cell shape and organelle distribution in astrocytes treated for 2h with vehicle, EGF, Yoda-1 or Yoda-1 together with Gefinitinib (Fig. 4E). Both Yoda and EGF decreased the size of the astrocytes (possibly indicating a contraction of the actin cytoskeleton), an effect that was blocked by the co-treatment with Gefinitib (Fig. 4E-F) In contrast, EGF treatment alone induced a significant increase in astrocyte motility, reflected by increased net and total displacement (Fig. 4F–G). Yoda-1 treatment additionally induced marked reorganization of the endoplasmic reticulum, characterized by increased ER size, which was reversed upon EGFR inhibition with gefitinib (Fig. 4H).

Collectively, these findings indicate that Piezo1 opening is sufficient to trigger EGFR phosphorylation, signaling and the related restructuring of the cellular architecture and these effects are dependent on the kinase activity of EGFR.

### Piezo1 activation in astrocytes induces pro-inflammatory cytokines through EGFR

The clustering analysis of the porcine in-vivo expression data pointed toward a mechanistic link among ICP, Piezo and EGFR but also to the correlation between ICP and a number of inflammation-related molecules such as CCL-2 and TIMP-1. We explored in human astrocytes the mechanistic association between these two aspects of the response to ICP. We set out to verify if Piezo1 opening is sufficient to induce pro-inflammatory cytokines and the eventual involvement of EGFR in this process. Therefore, we monitored the expression of CCL2, of IL-6 and IL-8 upon astrocytes treatment with EGF (2h and 48h), Yoda (2h, with 48h treatment being lethal to astrocytes) and Yoda-1 in combination with Gefitinib (2h). Both short- and long-term EGF treatment, as well as Yoda-1 treatment, resulted in the robust increase in the expression of all three cytokines. Notably, this induction was reversed by concomitant gefitinib treatment, indicating that cytokine upregulation is dependent on EGFR kinase activity.

Similarly, EGF and Yoda induced the upregulation of TIMP1; however, this effect was not reversed by EGFR kinase inhibition (Fig. 5D), suggesting the presence of a redundant signaling pathways not requiring EGFR. Interestingly, although neither Yoda nor EGF affected AQP4 or S001B mRNA, Gefitinib + Yoda treatment substantially upregulated both genes, implying that baseline EGFR activity is necessary to suppress AQP4 and S100B (Fig. 5E–F). In order to confirm the identity of the downstream signaling cascades, we employed a pharmacological ERK inhibitor (Fig. G–L). ERK inhibition suppressed the induction of the pro-inflammatory cytokines IL-6, IL-8, and CCL2 upon Piezo1 activation. In contrast, combined treatment with Yoda-1 and ERK inhibitor increased the expression of AQP4 and S100B, whereas TIMP1 regulation followed a pattern similar to that observed in the gefitinib experiments and was not affected by ERK blockade.

**Figure 5.**
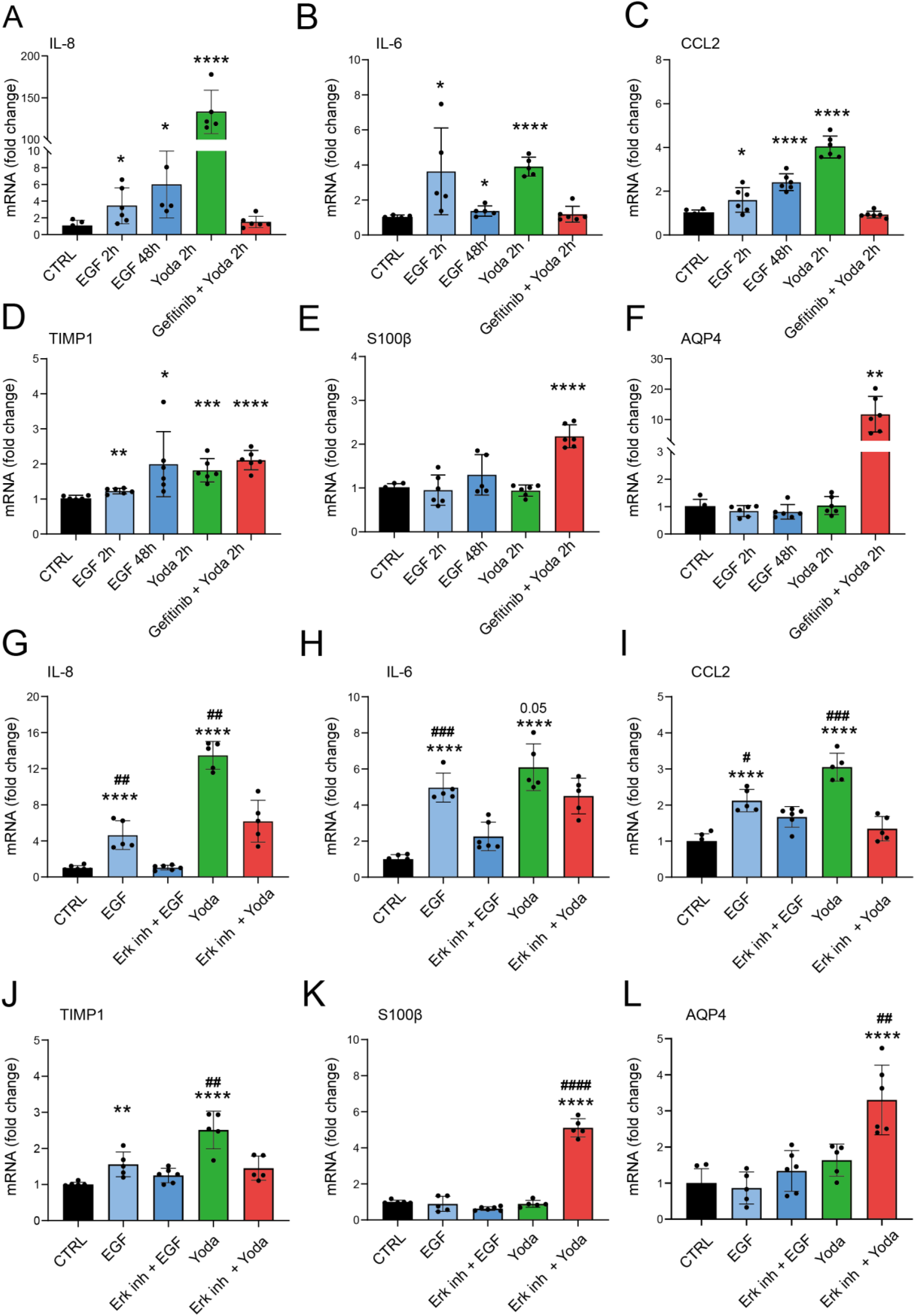
Piezo1–EGFR signaling drives pro-inflammatory cytokine expression and is suppressed by EGFR inhibition. (A–C) qPCR analysis of pro-inflammatory cytokines (CCL2, IL-6, IL-8) in human astrocytes following EGF, Yoda-1, or Yoda-1 plus gefitinib treatment. (D-F) Expression of TIMP1, AQP4 and S100B following EGF, Yoda-1, or Yoda-1 plus gefitinib treatment.(G–I) qPCR analysis of pro-inflammatory cytokines (CCL2, IL-6, IL-8) in human astrocytes following EGF,EGF plus ErK inhibitor, Yoda-1, or Yoda-1 plus ErK inhibitor treatment.(J-L) Expression of TIMP1, AQP4 and S100B following EGF,EGF plus ErK inhibitor, Yoda-1, or Yoda-1 plus ErK inhibitor treatment. Data are presented as fold change relative to CTRL. * indicates statistical significance compared with the CTRL group; # indicates statistical significance compared with the corresponding inhibitor-treated group. Data are presented as fold change relative to CTRL. Unpaired t-test vs CTRL was used to compare the groups Each group includes n = 6 independent samples. Data are shown as mean ± SD. *p < 0.05, **p < 0.01, ****p < 0.0001 vs. CTRL; #p < 0.05, ##p < 0.01, ###p < 0.001, ####p < 0.0001 vs. the corresponding inhibitor-treated group.

Together, these results indicate that Piezo1 activation is sufficient to induce pro-inflammatory cytokines in astrocytes through a pathway that involves the EGFR signaling.

### EGFR phosphorylation correlates with CCL2 expression, ICP and survival in the ASDH in vivo model

Finally, we investigated whether EGFR activation in the ASDH porcine model was correlated to the induction of neuroinflammation, as suggested by the human-astrocytes in vitro data. At the tissue level, robust ERK phosphorylation was detected in the brain parenchyma at 48h ASDH (Fig. 6A-B), confirming the preservation of phosphoepitopes. When probed for total EGFR and phospho-EGFR (Y1068), cortical samples from ASDH displayed a decrease in total EGFR compared to naive, but at the same time an increase in phosphorylated EGFR, indicating an increase in the fraction of activated EGFR (Fig. C-E)-and coherent with the degradation of EGFR upon persistent stimulation. We explored the levels of two cogntate ligands of EGFR, EGF itself and Neuregulin-1, finding both of them significantly elevated and supporting the ligand-induced EGFR signaling (Fig. 6F-H).

**Fig. 6.**
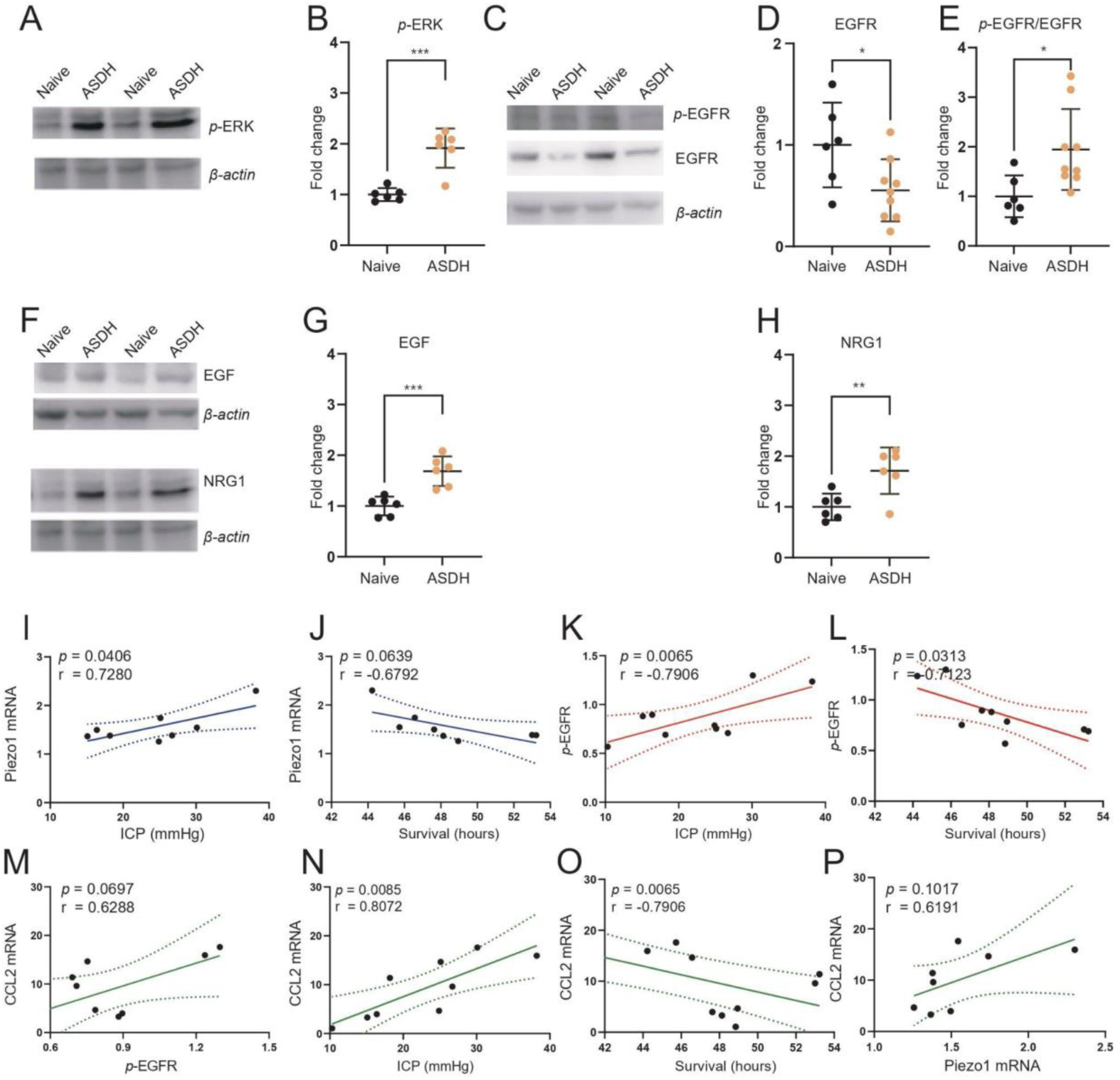
Piezo1–EGFR signaling correlates with intracranial pressure, inflammation, and outcome in the porcine ASDH model. (A–B) Immunoblot analysis and quantification of ERK phosphorylation in porcine brain parenchyma. Unpaired t-test was performed. Naïve animals n = 6; ASDH animals n = 6. ***p < 0.001. (C–E) Immunoblot analysis and quantification of EGFR phosphorylation and total EGFR levels in porcine brain tissue. Unpaired t-test was performed. Naïve animals n = 6; ASDH animals n = 9. *p < 0.05 (F–H) Immunoblot analysis and quantification of EGFR ligands (EGF and NRG1) expression in injured versus naïve cortex. Unpaired t-test was performed. Naïve animals n = 6; ASDH animals n = 6. ** p < 0.01, ***p < 0.001 (I–J) Correlation analyses between Piezo1 mRNA expression, intracranial pressure (ICP), and survival time. Pearson correlation was performed (K–L) Correlation of p-EGFR levels with ICP and survival time. Pearson correlation was performed (M–P) Correlation analyses of CCL2 mRNA expression with p-EGFR, ICP, survival, and Piezo1 mRNA expression. Pearson correlation was performed

**Figure 7.**
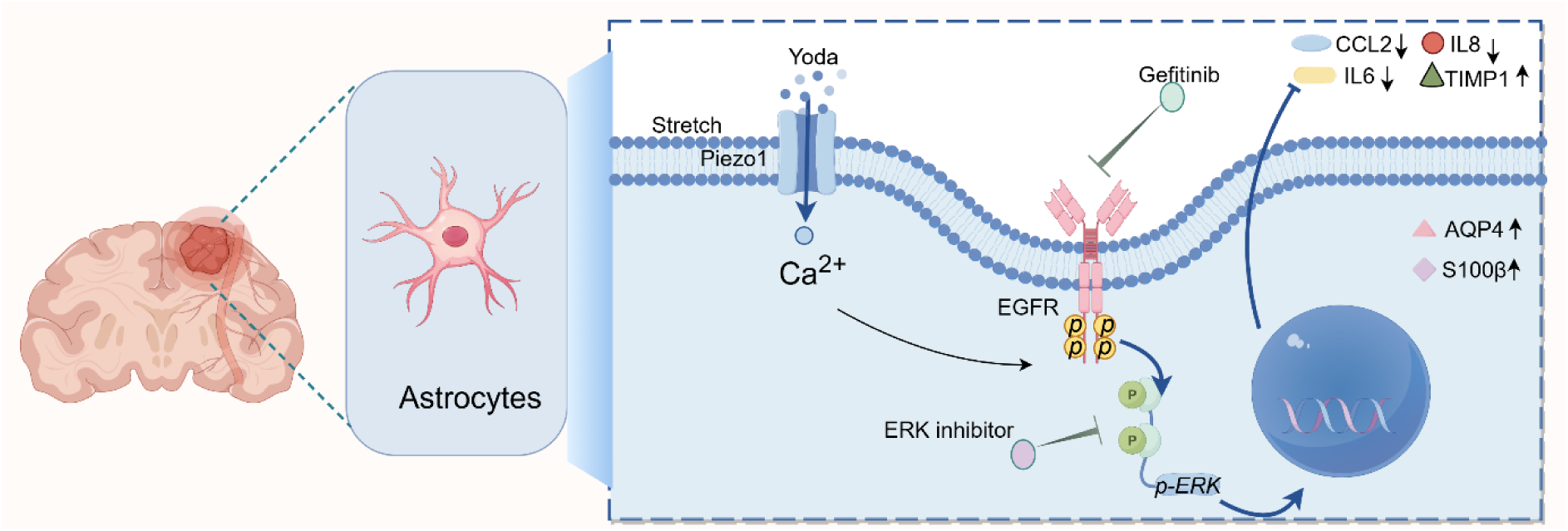
Elevated ICP and mechanical stress activate astrocytic Piezo1 mechanosensitive ion channels, triggering Ca²⁺ influx and site-specific EGFR phosphorylation This induces EGFR internalization and downstream ERK signaling activation, promoting the release of pro-inflammatory cytokines (CCL2, IL-6, IL-8) and thereby driving neuroinflammation and exacerbating cerebral edema.Pharmacological inhibition of EGFR or ERK suppresses the Piezo1-EGFR-ERK axis, attenuates inflammatory signaling, and markedly upregulates aquaporin-4 and S100β expression, shifting astrocytes toward a phenotype associated with water handling and edema containment.

We further examined if the Piezo1-EGFR-CCL2 axis displayed any correlation with clinico-physiological readouts such as ICP, CPP or survival. Piezo1 mRNA expression correlated positively with ICP and negatively with survival (Fig. 6I–J). Furthermore, phosphoEGFR levels were positively correlated with ICP, but inversely correlated with survival; in agreement with the in vitro data, pEGFR levels were directly correlated with CCL2 expression (Fig. 6 K-M). In fact, CCL2 expression itself directly correlated with ICP and Piezo1 expression and inversely correlated with survival (Fig. 6N-P).

Taken together, the in vivo data support the association between EGFR phosphorylation and ICP on one side and survival on the other.

## Discussion

Here we have provided convergent evidence, from an in vivo porcine ASDH model and in vitro human astrocytes, for a ICP-Piezo1-EGFR-CCL2 axis connecting intracranial hypertension associated with the subdural bleeding to astrocyte reactivity and neuroinflammation set in motion by Piezo-1 mechanosensation and mediated by EGFR phosphorylation. This axis appears to be highly relevant in shaping the acute prognosis, as its activity highly correlates with survival.

The first component of the axis is the mechanosensitive response to ASDH. We detect the increased expression of several mechanosensitive channels, including Piezo1, Piezo2 and TRPC3, which implicates the hypersensitivity to pressure and mechanical stress occurring upon ICP increase. Interestingly, a similar upregulation of pressure sensors has been identified in intracerebral hemorrhage models [36] and in sub-arachnoid hemorrhage [37], indicating that this may be a general mechanism set in motion to trigger compensatory mechanisms to bring the ICP down to the physiological range. To this respect, different mechanosensors may be involved in different aspects and cellular sites of this coordinated adjustment: whereas TRPC1,4 and 5 display an extensive neuronal expression [38], TRPC3 has been associated to pericytes response to ICP [37]. Conversely, Piezo1 and Piezo2 have a broad expression pattern and have been reported to be associated, among other effects, with neuronal responses to ICP [39, 40]. Here we have used a cross-correlation analysis of targeted gene expression to point toward the association of Piezo1 expression and ICP responses in astrocytes. Astrocytic roles for Piezo1 have been previously reported in terms of induction of reactive phenotypes [41, 42] and regulation of glymphatic flux [43]. Our findings implicate Piezo1 in astrocyte response to ICP in ASDH.

The second component of the axis is constituted by the link between Piezo1 and RTK signaling. EGFR emerges from the gene-expression correlation analysis, together with VEGFR1 and TrkB, as the most correlated with ICP. Of note, the pattern of up-regulation for RTKs and their ligands upon ASDH is actually broader and includes, among others, Axl, FGFRs, Erb and HGFR, coherently with the observation of their role in neuronal responses and neuroinflammation upon injury [17, 44-46]. We prioritized EGFR because genetic evidence (in the oncology field) suggests its prominent role in driving astrocytes proliferation [30, 47] and EGFR inhibitors are available for human use [48]. Interestingly, EGFR phosphorylation has been previously correlated, in non-cerebral systems, with changes in cell-shape [49] rather than direct physical interactions. Our phosphoepitope screening suggests multiple similarities between EGF-induced and Piezo1-induced EGFR phosphorylation (including the rapid internalization of the EGFR, implicating its activated signaling;[50], although the two phosphorylation patterns do not fully overlap. Notably, a similar Piezo1-EGFR cross-activation has been previously observed in HeLa cells and lung tissue [51]; however, in this context the signaling through EGFR was not sensitive to EGFR tyrosine kinase blockade but rather to Src. Interestingly, an alternative non-cell-autonomous cross-talk pathway has been described in a liver regeneration model, in which Piezo1 activation in endothelial cells enhances the release of EGFR ligands and drives EGFR signaling, sensitive to Gefitinib, in hepatocytes [52]. Thus, at least two mechanisms of EGFR activation by Piezo1 may be active (non-mutually exclusive and possibly redundant) in our system: one in which Piezo triggers the increased availability of EGFR ligands and activates EGFR-kinase dependent signaling, and another in which Piezo1 triggers the phosphorylation of EGFR by Src, with the downstream signaling being EGFR-kinase independent. In our system, Gefitinib (selected because of its translational value) obtains a consistent suppression of Piezo1-triggered effects suggesting the requirement for the receptor kinase activity; although Gefitinib is not considered a Src inhibitor, cross-reactivity has been suggested [52].

The third component of the axis is constituted by the response of astrocytes to EGFR. The proliferative response to EGFR and the transition of astrocytes to a reactive phenotype upon exposure to EGFR ligands have been previously described [53-55].

Our data point toward a direct involvement of EGFR in driving a local neuroinflammatory state sustained by astrocytic release of pro-inflammatory, chemotactic mediators: CCL2 is among the genes that closely cluster with Piezo1 and EGFR expression in the porcine in vivo samples, and CCL2, IL-6 and IL-8 are upregulated either by EGF either by Yoda1 treatment in human astrocytes (an effect negated by Gefitinib co-treatment). In line with our observation, mutant EGFR was shown to contribute to macrophage recruitment in glioblastoma through CCL2 [56]. A similar effect of EGFR signaling was reported in breast cancer models, where EGFR induced CCL2 and IL-8 through the EGR1 transcription factor [57]. Furthermore, a cross-talk between EGFR and NF-kB through p38k has been shown to contribute to cytokine secretion and macrophage recruitment upon spinal cord damage [58]. Our findings confirm this to be a generalized effect of EGFR, driving proliferation and local inflammation at the same time. and are in general agreement with the previous observations of EGFR mediating some of the pro-inflammatory effects of increased hydrostatic pressure (e.g. in the optic nerve astrocytes [59]).

Finally, our in vivo data support the model in which the activation of the Piezo1-EGFR axis is relevant in prognostic terms, since the phosphorylation of EGFR is inversely correlated with survival and directly correlated with CCL2 expression. The finding, being based on a single datapoint, does not lend itself to the establishment of a linear causality relationships: on one side, ICP increase may be responsible for triggering EGFR phosphorylation, and simultaneously the higher the ICP, the shorter the survival; on the other side, higher EGFR signaling may induce a stronger reactive astrocyte state, reducing their ability to manage water fluxes and to compensate for the intracranial hypertension, further complicating the ICP increase. These two events may occur at the same time in a self-amplifying vicious circle: once the increased expression activates Piezo1-EGFR, the EGFR signaling moves astrocytes toward a reactive state in which they become less able to enable the adjustment to increased ICP, leading to a permanent or worsening increase in ICP. In this direction, EGFR inhibitors may be useful to break this cycle; in fact, EGFR inhibitors have proven to be beneficial in preclinical models of brain or spinal cord damage [58, 60].

The proposed model must be interpreted within the limitations of the present study, in particular the difficulties to establish direct causality links using cross-sectional data (from the in vivo model). In the same line of thought, the present study was designed to stop at 48h of follow-up (because of logistical and methodological consideration), therefore remains unclear if the Piezo1-EGFR axis remains relevant at earlier or later timepoints, i.e. the therapeutic window for EGFR-targeted interventions remains to be determined. A direct interventional study (with the EGFR inhibitor) will be better suited to establish the relevance of the Piezo1-EGFR axis in therapeutic terms.

## Supporting information

Supplementary animal procedure

## Acknowledgement

This work is supported by the Deutsche Forschungsgemeinschaft (DFG) in the context of the Collaborative Research Center SFB1149 “Danger Response, Disturbance Factors and Regenerative Potential after Acute Trauma” (Grant no. 251293561 to FR;PR). F.R. is also supported by the Bundesministerium für Bildung und Forschung (BMBF) under the DC4MND project (part of the JPND initiative; Grant no. BMBF-01ED2301) Z.Z is supported by the China Scholarship Council (Personal certificate No.202308320129).

## Authors’ contributions

Conceptualization. F.R., M.P.,T.K., P.R.; data curation Z.Z M.P.; formal analysis F.R. Z.Z. M.P.; funding acquisition and investigation F.R. P.R. methodology A.H. Z.Z. F.S. F.oH. T.M. B.O. M.G. S.K. E.C. F.M. M.P.,supervision F.R. M.P. T.K.; writing—original draft M.P. F.R. P.R., writing—review & editing F.R. P.R. T.K. M.P. All authors have read and agreed to the published version of the manuscript.

---

**Fig. S1.**
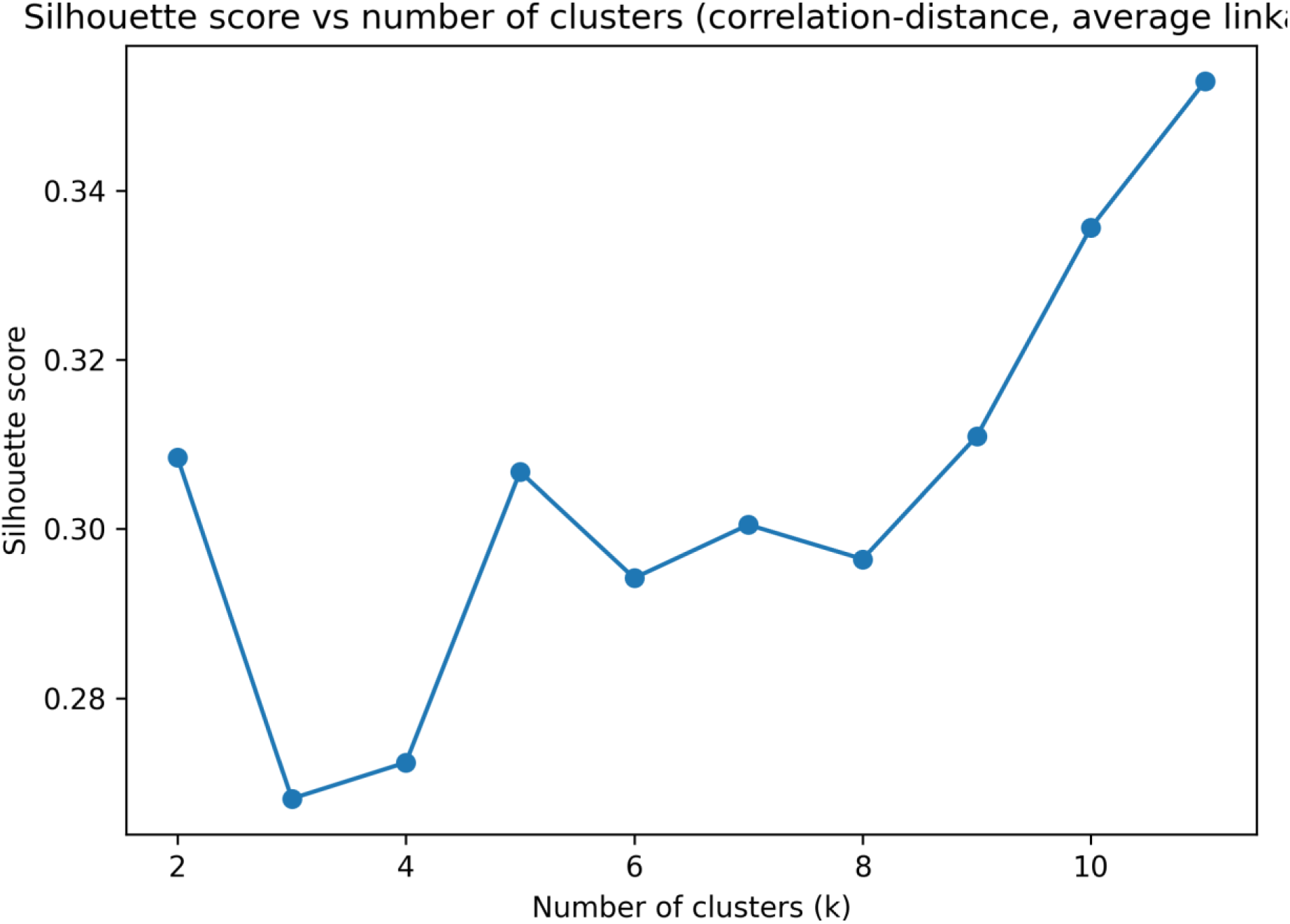
Silhouette scores for hierarchical clustering using correlation distance and average linkage across different numbers of clusters (k = 2–11).

**Fig. S2.**
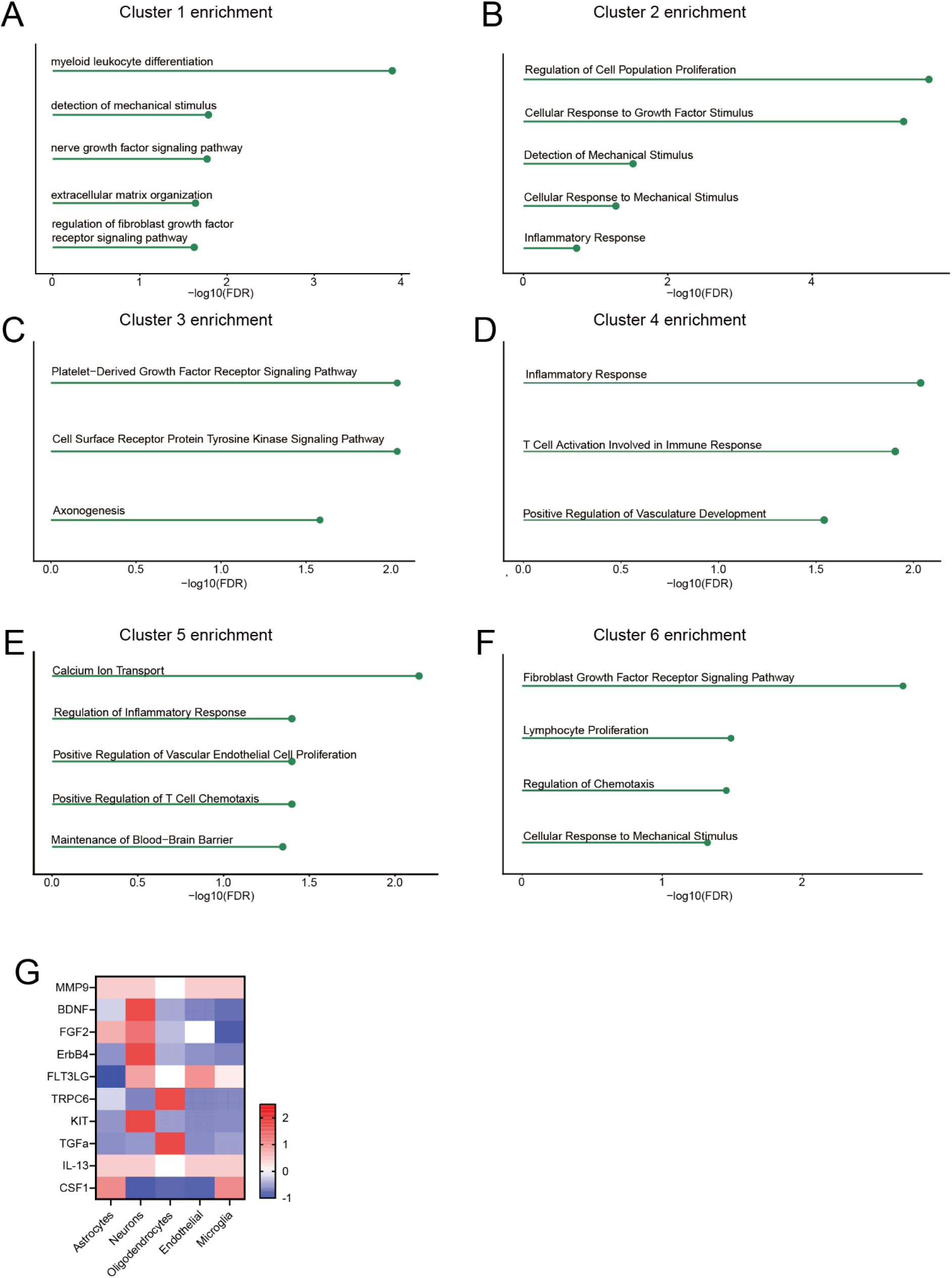
Gene ontology enrichment and cell-type–specific expression patterns. Gene Ontology (GO) enrichment analysis was performed for six gene expression clusters identified in the dataset. (A–F) Dot plots show the top enriched biological process terms for Cluster 1 to Cluster 6, respectively. The x-axis represents enrichment significance expressed as −log10(false discovery rate, FDR). Each dot corresponds to one enriched GO term. (G) Heatmap displaying expression levels of selected genes in Cluster1 across major brain cell types, including astrocytes, neurons, oligodendrocytes, endothelial cells, and microglia。

**Fig. S3.**
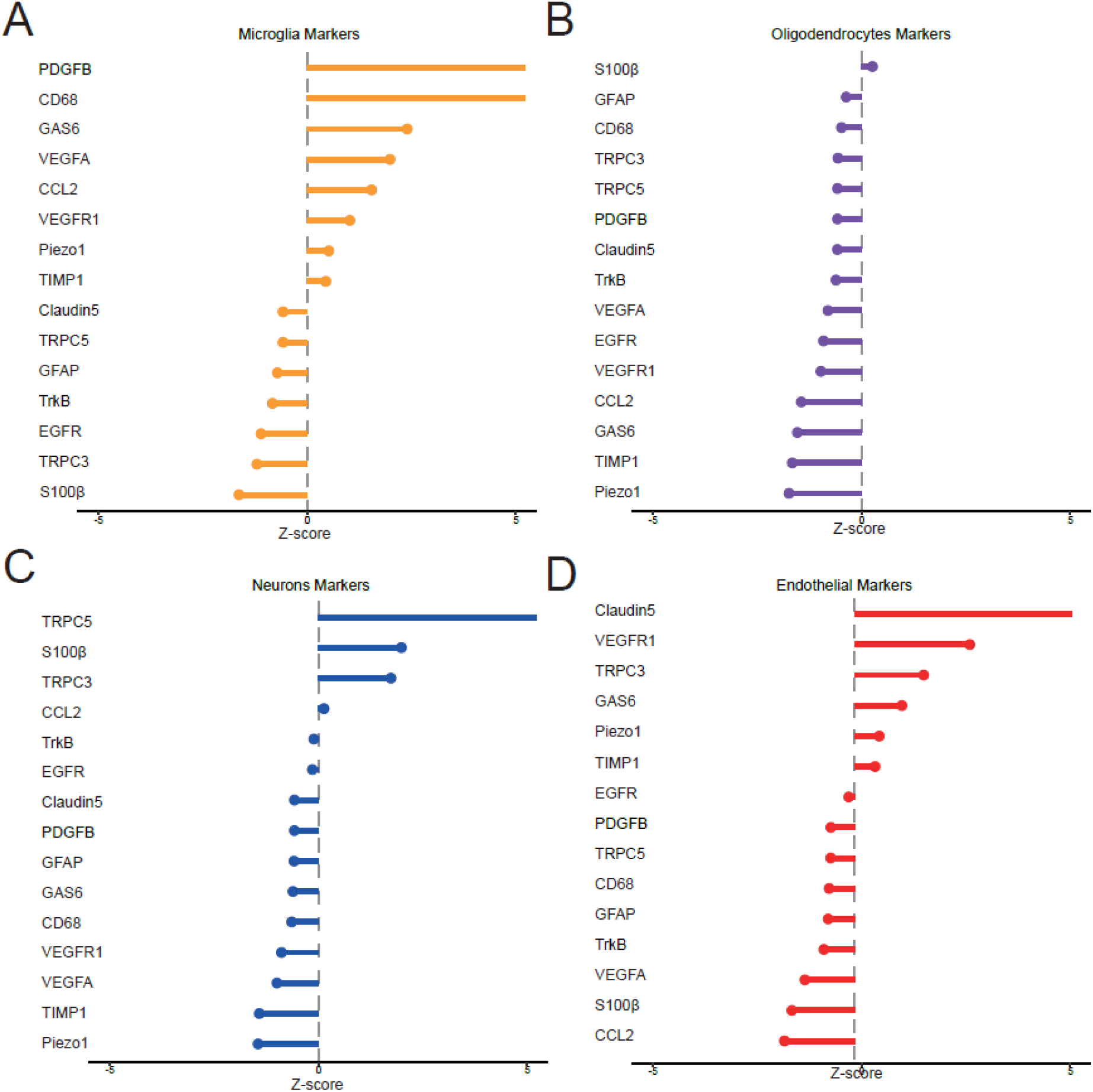
Standardized enrichment of cell-type–specific gene signatures across major brain cell populations. Cell-type deconvolution analysis showing the relative enrichment of selected marker gene signatures across major brain cell types. Z-scores indicate standardized enrichment levels for genes associated with (A) microglia, (B) oligodendrocytes, (C) neurons, and (D) endothelial cells

