## Supplementary animal procedure for "Intracranial hypertension drives astrocyte-mediated neuroinflammation through Piezo1-dependent EGFR activation"

**Supplementary material: Animal model and surgical procedures**

The porcine model of combined ASDH and hemorrhage has been previously reported and characterized (Datzmann et al., 2021; Datzmann et al., 2023; Kapapa et al., 2024). A total of 16 German Large White pigs (age 21 – 25 weeks, bodyweight 62 – 100 kg) of either sex were used for the present study. The animals were obtained from an agricultural production specified in breeding and rearing of piglets (Schweinezucht Kugler Gmbh & Co. Kg, Ostrach, Germany). The breeding facility is located approx. 100 km from our research facility, which helped to keep the transport of the animals as short as possible. The animals were kept at the breeding facility for up to 2 – 4 weeks before the experiments. After weaning, the piglets were kept in groups of 42 mixed-sex groups on polymer concrete, with access to a fodder rack with straw and were fed with fodder produced by the breeder. When the piglets reached approx. 30 kg of weight they were transferred to the subadult area where they were kept in groups of a maximum number of 190 animals divided in sections of a maximum number of 14 animals per group and separated by sex. Animals from different piglet groups were not mixed to prevent them from having to fight for a new hierarchy. The sub-adult area had a concrete slatted floor with access to a fodder rack with hay/straw/silage and animals were fed with fodder produced by the breeder. After transfer to our facility, the animals were kept separated by sex in separate boxes, but in contact to the neighboring boxes. They had regular run and were provided activity by the animal caretaker. They received the same fodder as before, provided by the breeder, additionally they had *ad libitum* access to hay and straw. Male and female animals were kept under the same conditions. At the breeding facility, regular health monitoring by the “Schweinegesundheitsdienst SGD” (pig health service) and the “Tierseuchenkasse” (animal plague office) was performed. After transfer to our facility, an incoming control comprised checking for normal breeding and behavior typical for pigs. In case of irregularities, the single animal was evaluated, and stool samples from all animals were checked for parasites. Neither the most dominant animals nor the runts of the litter were included in the experiments. Fights for the hierarchy were prevented by not mixing animals from different established social groups. Furthermore, the animals were kept in age-specific groups, and animals from different ages were not mixed. After the transport to our facility the animals were sheltered at Oberberghof, Ulm, Germany, until further use with an acclimatization period of at least two weeks. The animals were kept at a cycle of 12/12 hrs light/darkness and were monitored at least once daily, the housing temperature was set to 21 – 22°C with a humidity of 50 – 60 %.

Prior the experiment, animals had free access to water and received a caloric diet (Fresubin, Fresenius Kabi, Germany). Administration of a nutritional solution instead of a regular diet allows for feeding the animals and thus avoiding stress to hunger until the induction of anesthesia while at the same time the potential danger of regurgitation of solid food particles and their aspiration into the trachea is prevented during induction of anesthesia and prior to endotracheal intubation. Premedication consisted of intramuscular azaperone (5 mg/kg) and midazolam (1 – 2 mg/kg). After establishing venous access via an ear vein, anaesthesia was induced with propofol (1 – 2 mg/kg) and ketamine (1 mg/kg), followed by endotracheal intubation and controlled mechanical ventilation. Ventilator settings were tidal volume 8 mL/kg, respiratory rate 8 – 12 breaths/minute adapted to an arterial PCO_2_ (PaCO_2_) = 35 – 40 mmHg, inspiratory/expiratory ratio [I/E] = 1:1.5, fraction of inspiratory O_2_ (F_I_O_2_) = 0.3, positive end-expiratory pressure (PEEP) = 10 cmH_2_O to prevent formation of atelectasis. Anaesthesia was maintained with continuous intravenous propofol (10 – 20 mg/kg/hour), remifentanil (initial bolus 5 µg/kg, followed by 15 – 20 µg/kg/hour), and midazolam (0.5 – 1.0 mg/kg/hour). Maintenance fluids (10 mL/kg/hour) consisted of a balanced electrolyte solution (Jonosteril 1/1, Fresenius Kabi, Germany). A gastric tube was placed for stomach decompression and fluid drainage. After surgical exposure, a 9-F catheter sheath was inserted into the right femoral vein for placement of a 4-lumen venous catheter (Arrow International). Both femoral arteries were exposed for placement of a 4-F PiCCO catheter (PULSION Medical Systems) for continuous cardiac output, pulse pressure and stroke volume variation measurement, and a 10-F catheter for blood sampling and passive blood removal to induce hemorrhage. A catheter was placed in the urinary bladder via midline mini-laparotomy. Thereafter, the animal was turned into the prone position, the skull was exposed, and a craniotomy was drilled over the left and right parietal cortices. After exposing the dura, a small incision was made, and a catheter was inserted approximately 5 mm into the subdural space. On the contralateral side, a similar burr hole was placed. Afterward, microdialysis catheters and multimodal brain monitoring probes (Neurovent-PTO, Raumedic) were inserted about 10 – 15 mm into the brain parenchyma in both hemispheres. The multimodal probes were placed under visual control for intracranial pressure (ICP), brain tissue O_2_ partial pressure (P_bt_O_2_), and temperature measurements. The probes were calibrated before insertion according to the manufacturer’s specifications. After placement, all catheters were allowed to equilibrate for about 1 h, and recording was started when P_bt_O_2_ values were stable. At the end of the neurosurgical procedure, the burr holes were closed using bone wax, which also served for catheter, microdialysis and probe fixation. Bilateral neurosurgery was performed to avoid sham experiments, i.e., to comply with the 3R principle the hemisphere without ASDH served as the control for the hemisphere with ASDH. During surgery, hydroxyethyl starch 6%–130/0.42 (Vitafusal, Serumwerk, Bernburg) was used to maintain pulse pressure variation, with a maximum dose of 30 mL/kg according to the manufacturer’s specifications.

Prior to the start of the experimental protocol, in order to mimic the clinical situation where resuscitation measures are initiated only after trauma-and-hemorrhage, maintenance fluid infusion rate was reduced to 100 mL/h, ventilator settings were tidal volume 8 mL/kg, PEEP = 0 cmH_2_O, I/E ratio = 1:2, F_I_O_2_ = 0.21. Thereafter, 0.1 mL/kg body weight of autologous blood was injected over 15 min via the subdural catheter using a syringe pump. This subdural blood volume was injected because 10 % of the intracranial volume represents the threshold for supra-tentorial volume tolerance (Timaru-Kast et al., 2008). Immediately thereafter, hemorrhage was initiated by passive removal of blood over 30 min via the 10F-arterial catheter targeting 30 % of the calculated blood volume. The removed blood was stored at 4 – 8 °C in citrate-anticoagulated bags. Blood removal was slowed if necessary to maintain cerebral perfusion pressure (CPP), i.e., the difference between mean arterial pressure (MAP) and ICP, ≥ 50 mmHg. After 105 min of hemorrhage, i.e., 2 hours of combined ASDH and hemorrhage, resuscitation was initiated comprising re-transfusion of shed blood within 30 min, fluid resuscitation (10 mL/kg/h; reduced to 5 mL/kg/h if venous pressure > 16 mmHg), and continuous i.v. noradrenaline titrated to maintain MAP at pre-shock levels and CPP ≥ 75 mmHg over a maximum of 48 hours. Upon initiation of resuscitation, the baseline ventilator settings were resumed (tidal volume 8 mL/kg, respiratory rate 8 – 12 breaths/min to maintain PaCO_2_ of 35 – 40 mmHg, I/E ratio of 1:1.5, and PEEP 10 cm H_2_O). The F_i_O_2_ was adjusted to maintain normoxemia, i.e., a PaO_2_ = 80 – 120 mmHg). Temperature management aimed to achieve brain normothermia, i.e., external cooling with ice-cold waterfilled bags was applied if brain temperature reached 39 °C. Neurological status was assessed daily using a modified Glasgow Coma Scale for pigs.

After 48 hours of intensive care the pigs were euthanized with KCl after further anesthesia deepening. Animals were euthanized before the end of the programmed 48-hours period upon predefined humane endpoints if *(i)* CPP < 60 mmHg despite maximum vasopressor dose, limited to a heart rate of 160/min to prevent tachycardia-induced myocardial injury, and/or the presence of *(ii)* refractory acute respiratory failure with PaO_2_ < 60 mmHg at F_I_O_2_ = 1.0, and/or *(iii)* anuric acute kidney failure with KC > 6 mmol/L. Target PaO_2_ values were always achieved using 0.23 < F_I_O_2_ < 0.3, and kaliemia always remained below 6 mmol/L. Hence, all experiments terminated pre-schedule were due to sudden ICP increases and a refractory fall of CPP < 60 mmHg.

Immediately post mortem, the head of the pig was removed. A skin incision on the middle of the forehead of the pig was performed, and the skin and muscle tissue were removed from the skull. The top of the skull was removed with the help of a bone saw and chisel. The dura mater was carefully opened with a scalpel. Afterwards, the pig head was balanced on the snout to facilitate the severing of the cranial nerves, after which the brain fell out of the skull and was gently caught by hand. The removal of the brain was always performed by the same trained and experienced veterinarian (AH). The time between the death of the animal and brain sampling was consistent for all experiments (30 ± 5 min). Without delay, brain tissue was immersed in formalin for preservation.

Naïve control animals (n = 6) underwent anesthesia alone without any neurosurgical instrumentation and induction of ASDH and hemorrhage. following the completion of the surgical catheter palcements, adjustments were made to the ventilator settings to achieve an inspiratory/expiratory (I/E) ratio of 1:2, F_i_O_2_ set at 0.21, and zero end-expiratory pressure (0 cmH2O) in order to closely replicate physiological conditions. These control animals were euthanized after a total experiment duration of approximately 2 hours and 15 minutes (Münz et al., 2024).
