## Supplementary Table 1 for "Intracranial hypertension drives astrocyte-mediated neuroinflammation through Piezo1-dependent EGFR activation"

Primers for RT-qPCR Analysis

| Gene | Forward Primer (5'-3') | Reverse Primer（5'-3'） |
| --- | --- | --- |
| *GAPDH* | CCTCCCCGTTCGACAGAC | ATGCGGCCAAATCCGTT |
| *TNF-α* | AACCTCAGATAAGCCCGTCG | ATTGGCATACCCACTCTGCC |
| *MMP9* | TCTTCTGGCGTGTGAGTTCC | AGGAGGTCGAAGGTCACGTA |
| *VEGFA* | AAGAAAATCCCTGTGGGCCT | CGTTTAACTCAAGCTGCCTCG |
| *VEGFR2* | TGCCTACCTCACCTGTTTCC | TGTTCTGCAGATACTGACTTAGAGA |
| *IL-33* | CCTTCTACAGCAGTTTGGCTC | TCTGTGTTCAGAGAGTTGCAG |
| *CD68* | ATCCTGCTGCCTCTCATCATT | GATGCAGAAGGTAACAAGCACC |
| *IL-10* | ACCTTCCAGGATGACGACTC | CGGGAACCTTGGAGCAGATT |
| *HGF* | GCTCATCGCAATAAAAAGCAGC | TCTCAGTGGAGGAGATGCCT |
| *MET* | TGGCATCCTTGTGCTTCTGT | CGTGGACTTTACCAGTGCCT |
| *CCL5* | TTTTTCCTACCTCTCCCGCC | TCTTCTCTGGGTTGGCACAC |
| *VEGFR1* | CGGTCTGCTTCTCACAGGATCT | ACGTGCTGGGTGCCTTTTA |
| *AQP4* | GGCTGCAGGTTCCAAATGTAAA | ACAGAGGTCCACACTTCCTTGTA |
| *EGFR* | CACCACCTACCAGATGGACG | ACCACGTAGTTACGAGGGCA |
| *ZO-1* | AGAGGAAGCTGTGGGTAACG | AGGTTTTCCTTGGCTGACACT |
| *CLDN5* | CTCTGCTGGTTCGCCAACAT | CAGCTCGTACTTCTGCGACA |
| *IGF1R* | ACGAGTGGAGAAATCTGCGG | ATGTGGAGGTAGCCCTCGAT |
| *IGF2* | CGGCTTCTACTTCAGACTTCCA | AGCAGCACTCTTCCACGATG |
| *FLT3* | GCTCCTCTGATGGTTACCCG | ATCTCTTCCGTGCAGTTGGG |
| *ErbB4* | TGGTCCCTCAGGCTTTCAAC | GCTGTGTCCAATTTCACTCCTG |
| *FGFR1* | GGAGGCTACAAGGTCCGTTA | ATACTCGTTCTCCACGACGC |
| *FGFR2* | TTACAACTCGCCTCTCCTCC | TGCCCAGCGTCAGCTTATC |
| *FGFR3* | ACGTGCACAACCTCGACTAC | CCGAAGGACCAGACATCACT |
| *Tie2* | GGGATCTACTCAGTTTTGGAACA | TCCAGAAGCAATGCAGGTGA |
| *PDGFRA* | TCTTTTCCCTTGGTGGCACA | CCATCCGGTACCCACTCTTG |
| *PDGFA* | GAGATAGACTCCGTAGGGGC | CTTCTCGGATGCATGCTTGG |
| *ANGPT1* | ATATACTGAGAGGAGGGTGCAG | GCCCATGCTGAATCCGGTTA |
| *HIF1a* | ACCATGCCCCAGATTCAAGAT | TCACTGGGACTGTTAGGCTCA |
| *FGF2* | CACACAAACATGGAGATGCCA | ATTGGGGTGCAGGTACCAAA |
| *FLT3LG* | TACTGCTGGTCGAGAAAGGC | TGGTTCACTCCTGCCAACAA |
| *NRG1* | ACAGCCCCCGCCAATAAATA | ACTCTGCCTAGGTCTCTCGC |
| *BDNF* | ATAGAGTCTGGGGATTTCGGG | CACCTGGTGGAACTTTTCAGTC |
| *AXL* | CCGTGAAGACGATGAAGATTGC | TTCCTTCATGCAGACGGCTT |
| *GAS6* | GAATTGCAGCTCCGCTACCA | TCCTCAACAGAGATTGTCTGCC |
| *EFNB2* | GCCGGACATTCTGGGAACAA | GCCGGACATTCTGGGAACAA |
| *EFNB3* | GTGCCTCTCCCATCTCTAGG | CTGAGGGGAGTGGTTGGTAAG |
| *EPHA4* | CCAGATCTGTTCAGGGAGAAC | AATACTCACTTCCTCCCACCCT |
| *CSF1R* | CAACAGTACCGACCCTACGA | TATTGGCCAGCCAAACCCTT |
| *c-KIT* | GTGGTCAAAGGAAACGCTCG | CCCATAGGACCAGACATCGC |
| *TYRO3* | GAGAAAAGTTGGGGCCTGAAG | TGTATCTCAGGCTCCTCCATCC |
| *TIMP1* | AGGAGTTTCTCATAGCTGGACA | TTCCAGGGAGCCACAAAACT |
| *TIMP2* | TCCTCAGGGACACAATTTGGAC | TGGGGTTGCCGTAGATGTC |
| *S100B* | TACCAGCTGTTTCCTTAGGTCG | CCAGTAGGAAGCTTCTGTCGC |
| *TGFα* | AAAGGACTTCTTGAGGTCGGG | AGATGCAAAGGCCGCATAGG |
| *GFAP* | GATTGTAAATGGAGCCCCGC | TTGAGGTGGCCTTCTGACAC |
| *VCAM1* | AGTTGCTCCCAGGGATACGA | GGAGCTGGAAAGCCATCACT |
| *ICAM1* | GCTCCAAACTTATGTCCTGCC | ACAGTTCACAGAAACGGGTGT |
| *IL-1β* | CTCCAGCCAGTCTTCATTGTTC | TGTCACCGTAGTTAGCCATCA |
| *CCL2* | ATTCTCCAGTCACCTGCTGC | TGCTGGTGACTCTTCTGTAGC |
| *IFN-γ* | CCAGGCCATTCAAAGGAGCA | TCAGTTTCCCAGAGCTACCA |
| *IL-4* | TGTGCTTCGGCACATCTACA | CATGTTTGCCATGCTGCTCA |
| *IL-13* | AGATGGGCAAACGAAGTGGA | TGTGGGAATCTGCGGGTTTTA |
| *TrkB* | AGTTTGGCATGAAAGGTTTTGTT | CAACCAACAAGCACCACAGC |
| *GJA1* | AGACAGGTCTGAGTGCCTGA | GCCCGGACACTACTCTTTCC |
| *PIEZO1* | GGACATCTACGCCAACGTCT | CCCATGCCGTACTTGACGAT |
| *PIEZO2* | GAGCTGGTGGTCTTCAACGA | ATCCCATAATGCCGTAGCCG |
| *TRPC1* | GTGCTTGGGAGAAATGCTGTT | CGCTCCATGAGTTTCTGACAA |
| *TRPC3* | TCCTGCTCTGTAGAGGGACTT | TTGTGTCAGTCTGCGGGAC |
| *TRPC4* | CGCTGTAACTGCGTCGAATG | GCCAGGGCCTTGTAGATGTT |
| *TRPC5* | CTGGTCCGTATTCGGCCTTT | ATGTCCCAAACATGGTCGCT |
| *TRPC6* | GGCCCTGAAAGTCGTGGTTA | GCATCCTCCGTCATGATCCC |
| *TRPC7* | TCGCCAGACTAGCCAACATC | CGCCCACCACAAAATCCTTG |
| *Human* |  |  |
| *TIMP1* | TTTTGTGGCTCCCTGGAACA | TGTGCATTCCTCACAGCCAA |
| *AQP4* | GCAGAAGCATTCTTTCTTGGTGT | CCTGCCATCTTCGGGTACTG |
| *CCL2* | AGAGGCTGAGACTAACCCAGA | TTTCATGCTGGAGGCGAGAG |
| *S100B* | TGAGACAAGGAAGAGCCCCTA | TGGAAAACGTCGATGAGGGC |
| *IL-6* | TCCGGGAACGAAAGAGAAGC | GAGAAGGCAACTGGACCGAA |
| *IL-10* | TCTGTCTGGACCCCAAGGAA | ATGAATTCTCAGCCCTCTTCAA |
