## Supplementary table 2 for "Intracranial hypertension drives astrocyte-mediated neuroinflammation through Piezo1-dependent EGFR activation"

| Antibody | Company | Cat. Nr. | Concentration |
|---|---|---|---|
| Erk (P-p44/42) MAPK (T202/Y204) | CST | 9101S | 1:1000 |
| Rabbit anti-Tie2 | My BioSource | MBS9611136 | 1:1000 |
| Rabbit anti-pTie2 (Ser1119) | My BioSource | MBS9614194 | 1:2000 |
| Rabbit anti-EGFR | My BioSource | MBS4511927 | 1:750 |
| Rabbit anti p-EGFR (Y1069) | Cohesion Bioscience | CPA3911 | 1:750 |
| Rabbit anti-VEGFR2 | My BioSource | MBS2028151 | 1:500 |
| Rabbit anti-pVEGFR2 | Cohesion Bioscience | CPA4047 | 1:750 |
| Goat anti-b actin | Santa Cruz Biotech | SC-1615-HRP | 1:1000 |
| Rabbit anti-NRG1 | Invitrogen | PA5-36057 | 1:1000 |
| Rabbit anti-EGF | My BioSource | MBS2006805 | 1μg/mL |
